# Thermodynamic properties and stability of HMGB1 complexes with linear polyelectrolytes elucidated by nano differential scanning fluorimetry

**DOI:** 10.64898/2026.08.27.747546

**Authors:** John Watson, Anna Klumpp, Eunjin Moon, Mariia Traviankina, Clemens Krage, Marten Kagelmacher, Marina Pigaleva

## Abstract

The High Mobility Group Box 1 (HMGB1) protein performs multiple essential functions in the body, ranging from DNA regulation to the activation and mediation of immune responses. However, HMGB1 has been also implicated in several pathological conditions, such as rheumatoid arthritis, sepsis, autoimmune diseases, tumors, and Alzheimer’s disease. As a result, HMGB1 is of increasing interest as a therapeutic target. Binding to heparin has been reported to inhibit HMGB1’s pathological activity during sepsis in clinical settings. In this work, we compare the interactions of HMGB1 with heparin and its’ synthetic analog linear polyglycerol sulfate (lPGS) from the viewpoint of stability and changes to association behavior. This analysis focuses on thermal stability, secondary-structure changes, and particle-size evolution using nano-differential scanning fluorimetry (nanoDSF), circular dichroism spectroscopy (CD), and dynamic light scattering (DLS).

## Introduction

HMGB1 is a multifunctional 215-amino acid residue protein that participates in both nuclear regulation and extracellular immune signaling [1,2]. Structurally, HMGB1 consists of two domains, the A box and B box, which possess a positive net charge at physiological pH [3,4] and are known to electrostatically bind to DNA, followed by a highly acidic C-terminal tail. The A and B boxes adopt predominantly α-helical folds [5,6], whereas the C-terminal tail is conformationally flexible and enriched with negatively charged residues. The tail modulates the accessibility and DNA-binding affinity of the HMG boxes, thereby contributing to HMGB1-dependent chromatin organization, transcription, replication, and DNA repair [4].

The biological function of HMGB1 strongly depends on its localization and redox state of three highly conserved cysteines (C23, 45 and 106) [7]. In the fully reduced form, which is commonly associated with the nuclear environment, HMGB1 contains three free thiols and binds nucleic acids as well as protein partners with relatively broad specificity. Upon its release into the extracellular space, fully reduced HMGB1 forms a heterocomplex with CXCL12. This complex activates the CXCR4 receptor and enhances CXCR4-mediated chemotaxis, thereby contributing to inflammatory cell recruitment [2,8]. Elevated extracellular HMGB1 levels have been implicated in several inflammatory and pathological conditions, including sepsis, rheumatoid arthritis, autoimmune diseases, cancers, and neurodegenerative disorders [1]. Consequently, targeting the pathological activity of HMGB1, without altering its essential intracellular functions, remains an important therapeutic and biophysical challenge.

One possible approach to modulating HMGB1’s activity is via inhibitory complexation with anionic macromolecules. This strategy is inspired by the natural affinity of HMGB1 for polyanionic binding partners such as DNA and by previous reports showing that sulfated glycosaminoglycans, including heparin, bind HMGB1 and interfere with HMGB1-mediated inflammatory signaling [9]. Heparin is commonly administered as an anticoagulant but also showed anti-inflammatory properties in the context of sepsis [10]. The interaction of heparin with HMGB1 has been suggested to contribute to anti-inflammatory effects [11,12]. However, heparin is structurally heterogeneous and contains a complex distribution of sulfate, carboxylate, hydroxyl, N-sulfate, and N-acetyl groups along a rigid polysaccharide backbone [13,14]. This heterogeneity complicates the identification of molecular parameters that control HMGB1 binding, conformational response, and association.

Synthetic polyelectrolytes (PEs) provide a useful platform to address these questions in a more controlled manner. lPGS, for example, represents a comparatively simple heparin analogue with a high degree of sulfation, a more flexible backbone and uniform charge distribution. Previous studies have shown that HMGB1 can interact with lPGS [15] However, a direct comparison of their effects with those of heparin on HMGB1 stability, refolding behavior, and association remains limited.

In this work, we investigate how the chemical nature and flexibility of anionic PEs influence HMGB1–polymer complex formation patterns and HMGB1 thermal stability. Using nanoDSF, together with complementary analysis of refolding and association behavior during complex formation and subsequent heating, we compare heparin with lPGS. This approach allows us to assess how charge effects, backbone flexibility, polymer nature and protein glycosylation reshape the thermal stability and oligomerization/controlled association propensity of HMGB1, providing insights into the physicochemical principles governing HMGB1–PE interactions.

## Experimental

All measurements were performed in a reducing HEPES buffer containing 25 mM HEPES and 20 mM dithiothreitol (DTT) at pH 7. DTT was included to maintain HMGB1 in its fully reduced state. The NaCl concentration was 0 mM (salt-free conditions) and 150 mM (physiological-salt conditions).

Recombinant human HMGB1 expressed in HEK293 cells (eukaryotic HMGB1, eu-HMGB1) was purchased from Sino Biological, catalogue number 10326-H08H1. Prior to use, the protein was buffer exchanged into the reducing HEPES buffer using 3 kDa Amicon centrifugal filters. Protein concentrations were determined by absorbance at 280 nm using a Thermo Scientific NanoDrop 2000 spectrophotometer. For comparison, recombinant human HMGB1 was expressed in *Escherichia coli* (*E. coli)* cells (bacterial HMGB1, bac-HMGB1) and purified accordingly [16].

SDS-PAGE was performed using a 4%-20% gradient Mini-PROTEAN® TGX™ Precast Protein Gel (Bio-Rad, catalog number 451096). Both eu-HMGB1 as well as the bac-HMGB1 were loaded onto the gel at 2, 20 and 40 μg respectively, by mixing with reducing 4x Laemmli Sample Buffer (abcr GmbH, catalog number AB351078) and subsequently heating to 95°C for 5 min. The PageRuler™ Prestained Protein Ladder (Thermo Scientific, catalog number 26616) was used as a reference. The gel was run at 20 mA for 15 min and subsequently at 40 mA for 25 min. The Periodic Acid Schiff Assay (PAS) was performed as previously described by Butnarasu [17]. For the Coomassie staining, PageBlue Protein Staining Solution (Thermo Scientific, catalog number 24620) was used for 1h followed up by destaining in MilliQ water overnight (o.n.). All images were acquired using a Chemidoc^**TM**^ MP imaging system (BioRad).

Fractionated heparin with an average molecular weight of 15 kDa (cat. number CAS 2608411) was obtained from Calbiochem and prepared as a 100 µM stock solution in Milli-Q water. lPGS (10 kDa, polydispersity index (PDI) 1.6, 80% sulfation) and hydroxyl-functionalized linear polyglycerol (lPGOH, 10 kDa, PDI 1.6) were synthesized according to the procedure reported by Nie et al.[18].

Thermal unfolding and association behavior were analyzed using a NanoTemper Prometheus Panta instrument. 10 µL of sample was loaded into Prometheus high-sensitivity capillaries (NanoTemper, catalog number PR-C006) and all measurements were performed in duplicates. Samples were heated from 25 to 75 °C and subsequently cooled back to 25 °C at a rate of 0.5 °C min□^1^. Intrinsic fluorescence was recorded at 100% excitation intensity. DLS was recorded simultaneously on the same instrument to monitor temperature-dependent changes in particle size and association. DLS measurements were acquired with 10 acquisitions per capillary, an acquisition time of 5000 ms, and 100% laser power. HMGB1– PE titrations were performed by mixing 20 µM of HMGB1 with varying concentrations of the polymers ranging from 0 to 150 mol%. nanoDSF and DLS data were processed using OriginPro. Melting temperatures were determined from the first derivative of the fluorescence melting curves.

Far-UV circular dichroism spectra were recorded using a JASCO CD spectrometer equipped with a Peltier temperature-control unit. Measurements were performed in a Hellma Analytics 110-01-40 high-precision quartz cuvette with a 1 mm path length. Spectra were collected from 190 to 250 nm with five accumulations per sample. For temperature-dependent CD measurements, HMGB1 was measured either alone or in the presence of equimolar heparin, or lPGS. Spectra were recorded sequentially at 20, 30, 40, 50, 60, and 75 °C, followed by cooling to 50 and 20 °C to assess structural refolding after heating. Once the target temperature had been reached in the cell holder using the Peltier element connected to the thermostat, the samples were allowed to thermally equilibrate at the target temperature for 5 min before each measurement.

## Results and discussion

### Thermal stability and refolding behavior of HMGB1 in the presence of sulfated polyelectrolytes

The thermal unfolding and refolding behavior of HMGB1 alone was first analyzed by nanoDSF (Fig. 1a). During heating from 25 to 75 °C, HMGB1 exhibited a single cooperative fluorescence transition, consistent with an apparent two-state unfolding process [19]. The main transition reflects changes in the local environment of the intrinsic tryptophan residues located in the A and B boxes, caused by exposure of these residues to the aqueous phase during unfolding [16]. The inflection point of this transition was assigned as the apparent melting temperature, T_m, app_.

**Figure 1.**
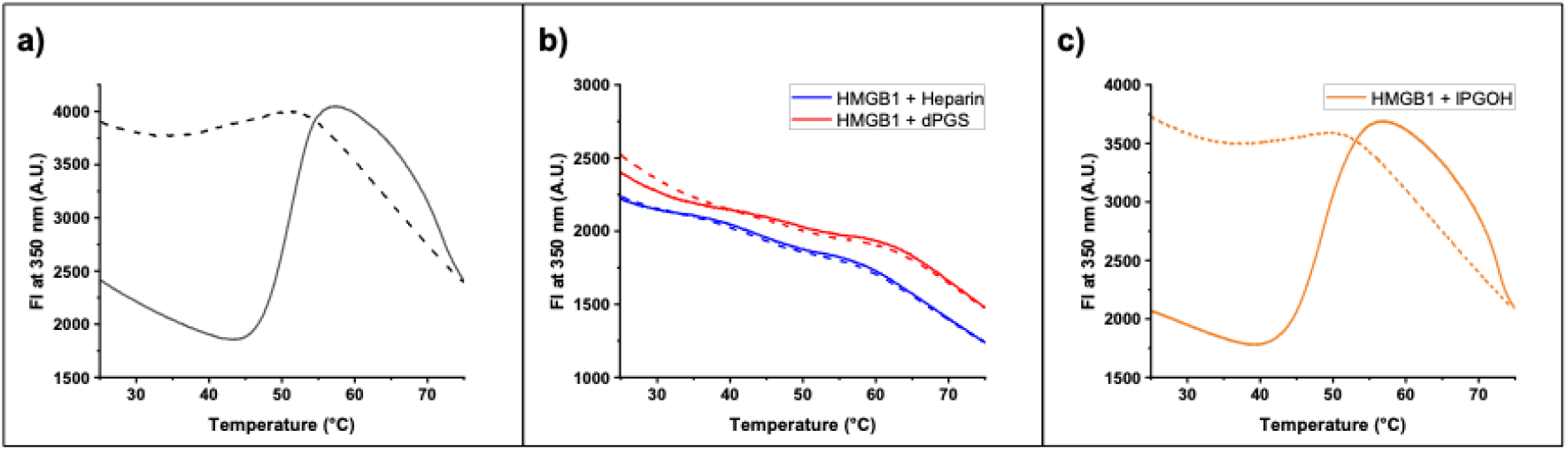
Thermal unfolding and refolding of HMGB1 monitored by nanoDSF. Thermal unfolding traces are shown as solid lines while refolding traces are indicated by dashed lines. Fluorescence intensity at 350 nm traces of HMGB1 during heating from 25 to 75 °C and subsequent cooling to 25 °C. Panels are arranged from left to right: a) HMGB1 in the absence of Pes, b) HMGB1 with 100 mol % heparin (blue line) and lPGS (red line), c) HMGB1 with 100 mol %lPGOH.

Upon subsequent cooling, the fluorescence signal did not retrace the heating curve, indicating that thermal unfolding of HMGB1 is only partially reversible under these conditions and that the tryptophan microenvironment does not fully return to the native state. This behavior is most likely caused by oligomerization of the protein to very large particles during thermal unfolding (see SI, Fig. S1). Because nanoDSF reports only local changes around the tryptophan residues, CD spectroscopy was used to assess the changes of the global secondary structure. The CD data confirmed that HMGB1 loses most of its α-helical secondary structure during heating and does not recover its initial spectrum upon cooling, demonstrating that the thermally unfolded protein cannot efficiently refold in buffer alone (Fig. 2a).

**Figure 2.**
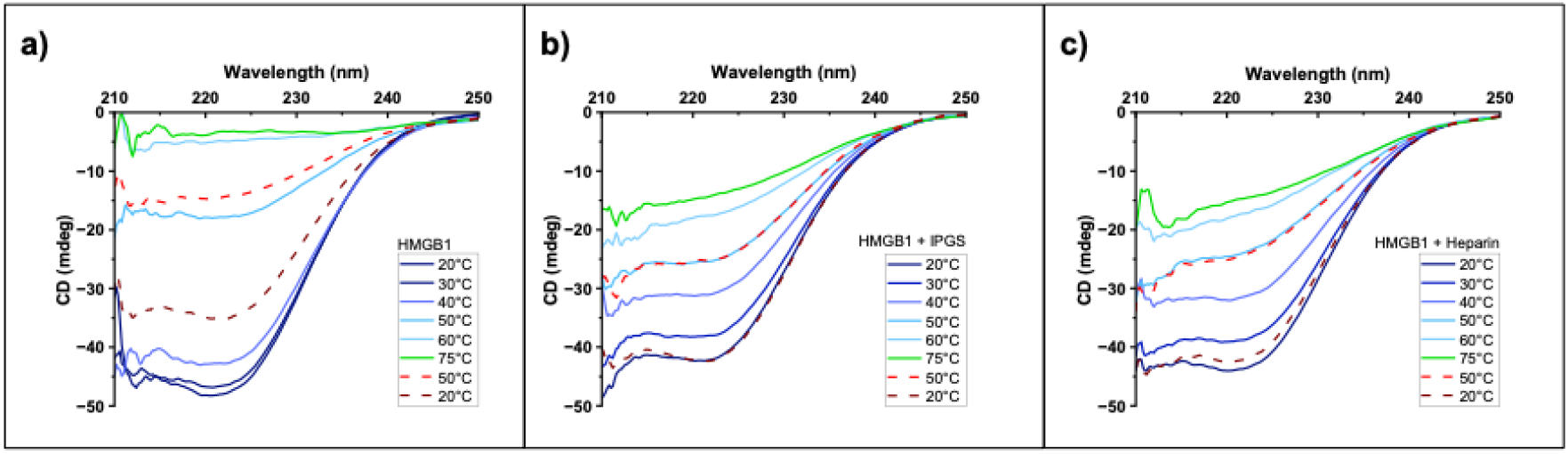
Circular dichroism spectra of HMGB1 and HMGB1–PE complexes recorded during heating to 75 °C, followed by cooling to 50 °C and subsequently to 20 °C. Spectra acquired during heating are shown as solid lines while spectra acquired during cooling are indicated by dashed lines. Panels are arranged from left to right: a) HMGB1 in the absence of PEs b) HMGB1 with 100 mol % heparin; c) HMGB1 with 100 mol % lPGS.

Under salt-free conditions, where electrostatic screening is minimized, the addition of equimolar heparin strongly altered the thermal response of HMGB1, indicating a pronounced contribution of electrostatic interactions between the polyanionic heparin chains and the protein (Fig. 1b, blue lines). In contrast to HMGB1 alone, the characteristic single unfolding transition was strongly suppressed, and the heating and cooling curves showed markedly reduced hysteresis. This indicates that heparin binding increases the apparent reversibility of the thermal response and protects HMGB1’s structure from irreversible unfolding. The binding of heparin apparently changes the unfolding pathway or stabilizes intermediate conformational states. One possible explanation is that the polyanionic heparin chain interacts with regions close to the tryptophan residues. Consequently the unfolding of the protein results in a smaller change of the local fluorescence environment. CD spectroscopy, in turn, showed that HMGB1 retains part of its secondary structure in the presence of heparin even at higher temperatures (Fig. 2b) and is able to fully refold upon subsequent cooling. Moreover, simultaneous DLS measurements during heating revealed that heparin stabilizes the hydrodynamic size of HMGB1 at (7.5 ± 1.3) nm (Fig. 3, a_3_) and prevents the oligomerization observed for HMGB1 alone in solution upon heating above 50 °C (resulting in the cumulant radius scattering from 100 to 1000 nm detected by DLS, see Figure S1).

**Figure 3.**
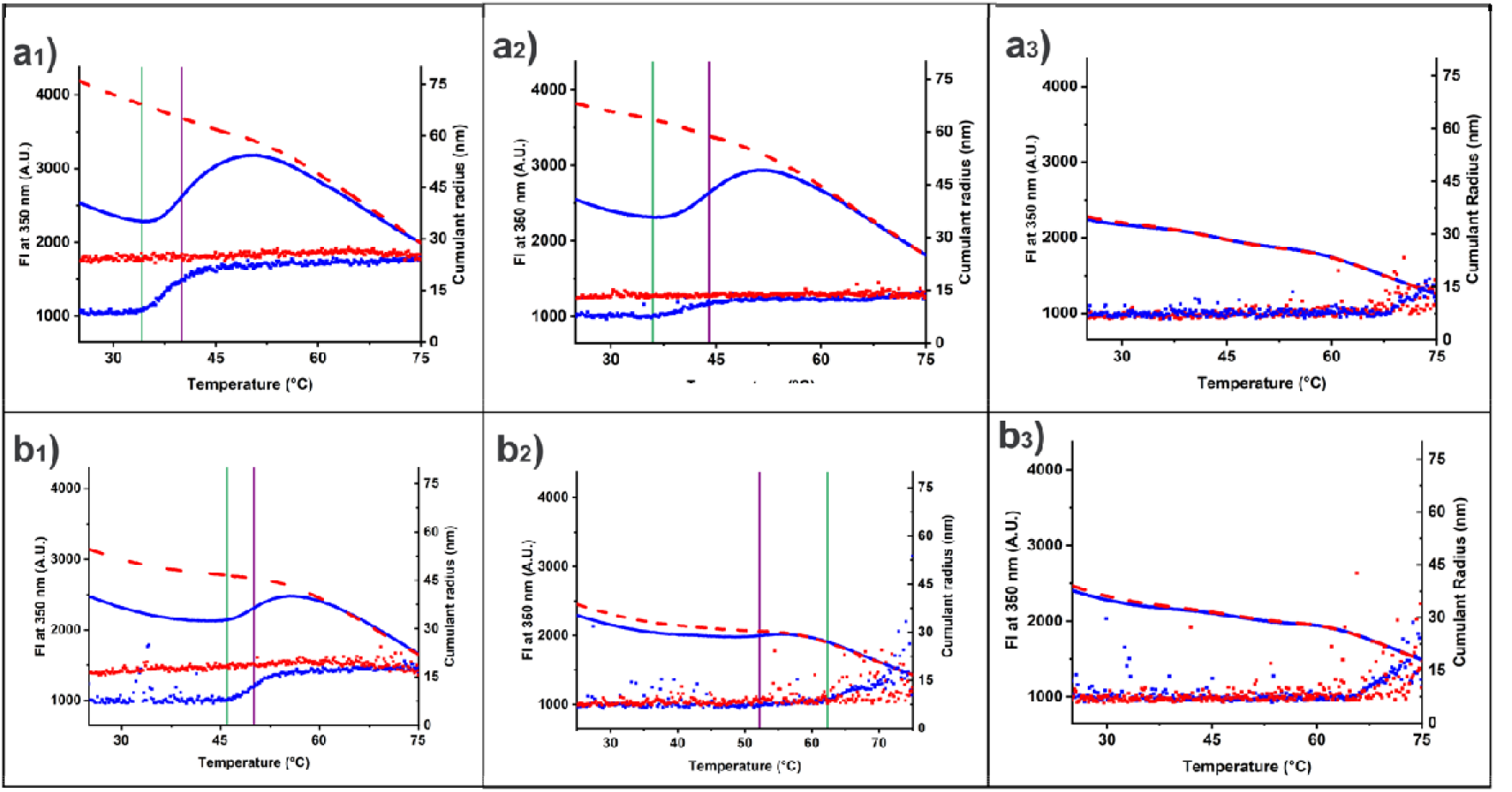
DLS and nanoDSF analysis of HMGB1 in the presence of PEs. Thermal unfolding traces are shown in blue and refolding traces in red; DLS data are presented together with the corresponding nanoDSF profiles above. Panels are arranged from upper left to bottom right: a_1_) HMGB1 with 30 mol% heparin; a_2_) HMGB1 with 50 mol% heparin; a_3_) HMGB1 with 100 mol% heparin; b_1_) HMGB1 with 30 mol% lPGS; b_2_) HMGB1 with 50 mol% lPGS; b_3_) HMGB1 with 100 mol% lPGS. Green vertical lines show the onset temperature of association detected by DLS, and purple vertical lines show the melting temperature.

A similar effect was observed by both nanoDSF and CD spectrometry for the synthetic heparin analog lPGS, stabilizing the hydrodynamic radius of HMGB1 at (8 ± 2.7) nm (Fig. 1b, red line; Fig. 2c). lPGS also suppressed the pronounced unfolding transition of HMGB1, enhanced the reversibility of the fluorescence signal upon cooling, prevented the temperature-induced loss of protein secondary structure, and enabled complete refolding to the initial conformation after thermal unfolding. This suggests that the stabilizing effect is not unique to the heparin structure but can also be induced by a synthetic sulfated polymer. Therefore, the interaction appears to be governed primarily by the presence of anionic sulfate groups rather than by the specific polysaccharide backbone.

To test whether the observed effects were primarily driven by electrostatic interactions between HMGB1 and the anionic PE or simply by polymer crowding or nonspecific steric shielding by macromolecular chains, HMGB1 was also measured in the presence of non-sulfated lPGOH (Fig. 1c). In contrast to heparin and lPGS, lPGOH did not suppress the unfolding transition and did not substantially improve refolding after cooling. The nanoDSF profile remained almost identical to that of HMGB1 alone. This control demonstrates that the polymer backbone itself is not sufficient to stabilize HMGB1. Instead, the effect requires sulfation and is therefore primarily driven by electrostatic interactions between HMGB1 and the anionic PE.

### Association behavior of HMGB1 in the presence of sulfated polyelectrolytes

While both heparin and lPGS are sulfated, linear polymers, strongly differ in their backbone flexibility, with heparin being a relatively rigid polysaccharide in contrast to the substantially more flexible lPGS (Kuhn’s length of (9 ± 0.1) nm vs. (0.87 ± 0.03) nm, respectively) [13,20]. To investigate whether these architectural differences influence the association behavior of HMGB1, DLS measurements were simultaneously acquired during the heating and cooling processes at different PE concentrations. To this end, it was observed that both heparin and lPGS can induce the formation of stable multimolecular HMGB1–PE assemblies during heating. For heparin, this behavior was observed over a broader concentration range, approximately 30–50 mol% (Fig. 3a_1_, a_2_), whereas for lPGS it was mainly observed around 30 mol% (Fig. 3b_1_). At 50 mol% polymer relative to HMGB1, lPGS suppressed the nanoDSF unfolding transition more efficiently than heparin (Fig. 3b_2_), indicating stronger stabilization of the tryptophan microenvironment at the same nominal polymer concentration. This narrower concentration window of added PE that results in distinct size assemblies observed by DLS with increase of the temperature for lPGS may reflect its higher conformational flexibility. A flexible sulfated chain can more efficiently adapt to the protein surface; screen exposed charged or hydrophobic regions and provide electrostatic repulsion between lPGS–HMGB1 complexes. As a result, lPGS may stabilize HMGB1 sufficiently to reduce the associate formation. In contrast, the more rigid heparin chain may require a narrower geometric match to the protein surface and may therefore stabilize HMGB1 in the found distinct-sized complexes state less efficiently at the same concentration.

The formation of these assemblies coincided with the thermal unfolding region of HMGB1, suggesting that partial exposure of hydrophobic protein regions may promote the distinct size nanometric association. In the case of heparin, this behavior is particularly relevant because association occurs within the range of physiological temperatures – starting from 34 °C. From this perspective, lPGS that stabilizes HMGB1 in its monomeric state until higher temperatures (46 °C) even under salt-free conditions, may represent a more favorable alternative.

### Influence of solvent ionic strength and protein glycosylation

In the following, we compared bac-HMGB1 with the eu-HMGB1 used before. Differences in glycosylation pattern between bacterial and eukaryotic expressed proteins have been observed for multiple fusion proteins [21]. Since HMGB1 contains four N-glycosylation sites (Asn 37, Asn 93 and Asn 134/135) we propose that the eu-HMGB1 shows a high degree of glycosylation [22]. Bacteria also tend to attach sugars to similar amino acids as eukaryotes, but in different patterns, less abundant and highly depended on the medium composition [21, 22]. To check for differences in glycosylation, a PAS assay of both eu-HMGB1 as well as the bac-HMGB1 variants was performed. PAS assay detects carbohydrate-containing molecules by oxidizing vicinal hydroxyl groups within sugar residues to aldehydes, which subsequently react with Schiff reagent to generate a characteristic magenta-colored product. The obtained results can be seen in Figure 4 for a) PAS staining, and b) control Coomassie staining. Both HMGB1 variants showed a PAS positive signal on the gel. It is known that *E*.*coli* cells tend to glycosylate fusion proteins: Geoghegan *et. al*. showed that the Histidine tag of fusion proteins can be spontaneously glycosylated with an α-*N*-6-Phosphogluconoylation [25]. Testing with lower amount of loaded protein (2 µg) revealed that the eu-HMGB1 still showed a visible band, whereas the bac-HMGB1 at the same concentration did not. Together with the results obtained for the 20 and 40 µg samples, these findings show that eu-HMGB1 exhibits a higher band intensity than bac-HMGB1, suggesting a higher degree of glycosylation.

**Figure 4.**
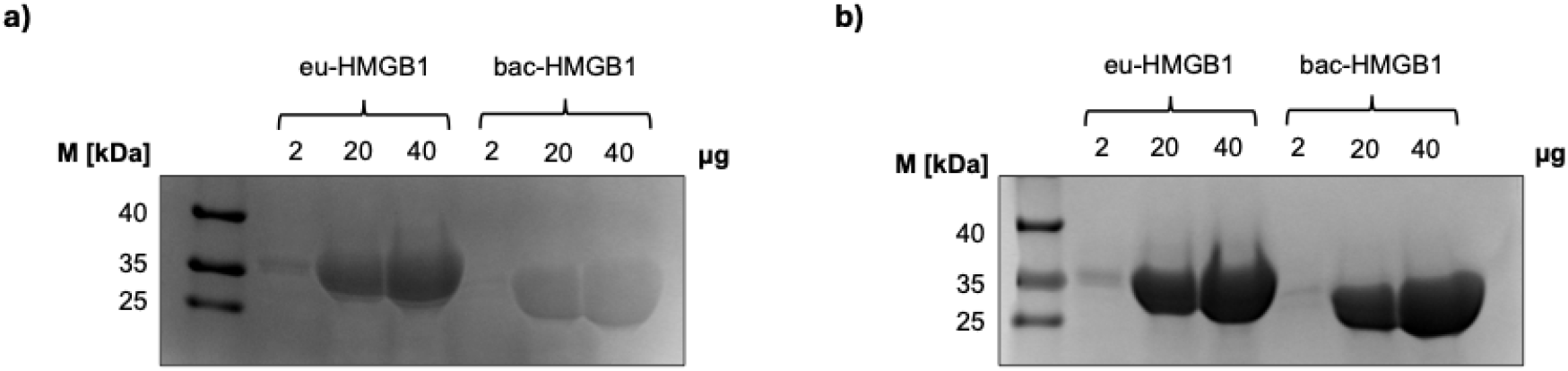
SDS-PAGE of eu- and bac-HMGB1. Samples were prepared at 2, 20 and 40 μg respectively. a) PAS, b) Coomassie.

To investigate whether this difference in glycosylation affects the unfolding behavior of HMGB1 and its association upon complexation with PEs, nanoDSF coupled with DLS was performed under the same salt-free conditions as described in the previous section. The association behavior of bac-HMGB1 was found to be largely comparable to that of eu-HMGB1 (Fig. 5a). In both cases, the cumulant radius started to increase at approximately the onset of unfolding, as detected by simultaneous nanoDSF measurements and the growth of the associates reached a plateau around the apparent melting temperature, T_m, app_. The final size of the established associates was slightly larger for the bac-HMGB1, reaching approximately 39 nm compared with around 26 nm for the eu-HMGB1. This may indicate that differences in sugar moieties on the protein surface contribute to the modulation of interprotein interactions, possibly by affecting the hydration shell, steric repulsion, and local surface accessibility of charged or hydrophobic regions, which could subsequently influence heparin-induced association during thermal unfolding. However, the overall similarity in the association profiles suggests that glycosylation is not the primary factor governing the temperature-triggered HMGB1–heparin association under these conditions. When the amount of heparin was increased to 100 mol% relative to the bac-HMGB1 concentration, the same general trend of refolding and size stabilization was observed (Fig. 5b). This further supports the conclusion that the formation of stable HMGB1–heparin associates is primarily driven by the interplay between protein unfolding and electrostatic complexation with the negatively charged heparin chains, rather than by the presence of glycosylation alone.

**Figure 5.**
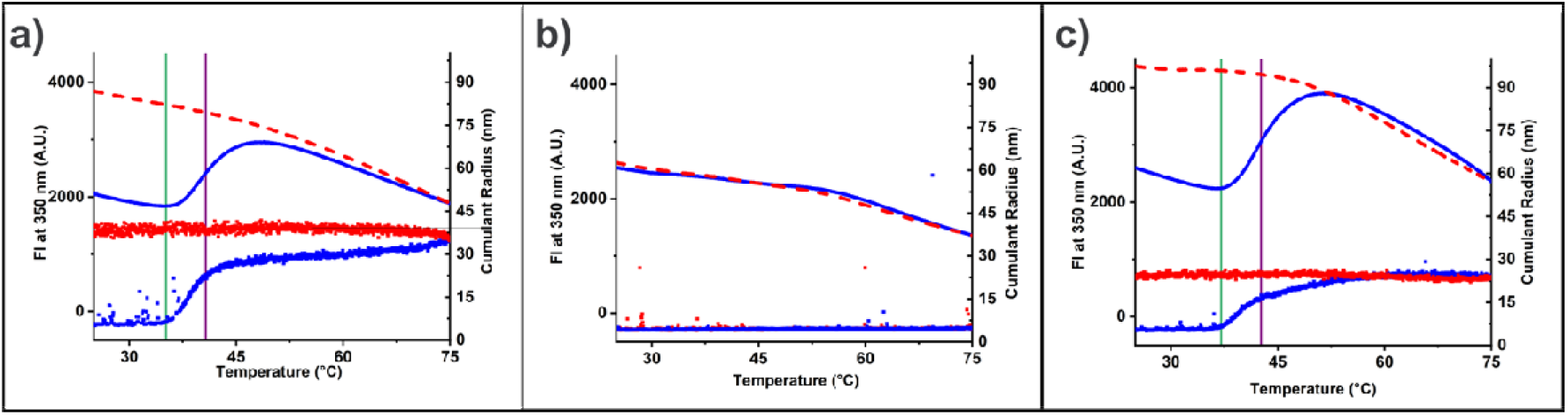
DLS and nanoDSF analysis of bac-HMGB1 in the presence of heparin. Thermal unfolding traces are shown in blue and refolding traces in red; DLS data are presented together with the corresponding nanoDSF profiles above. Panels are arranged from left to right: a) HMGB1 with 30 mol% heparin, salt-free conditions; b) HMGB1 with 100 mol% heparin, salt-free conditions; c) HMGB1 with 100 mol% heparin, physiological salt conditions.

To further probe these mechanisms at near-physiological ionic strength and to evaluate the influence of salt on HMGB1–heparin association, the salt concentration was increased to the physiological value of 150 mM. Under these conditions, 30 mol% added heparin was insufficient to stabilize the associates. However, at 100 mol% added heparin, bac-HMGB1 showed a trend (Fig. 5c) very similar to that observed at 30 mol% heparin under salt-free conditions. The onset of association was slightly shifted towards higher temperatures, from approximately 35 °C to 37 °C (see Table 1) but again coincided with the onset of unfolding measured by simultaneous nanoDSF. The initial, steeper increase in associate size proceeded approximately up to T_m,app_, followed by a slower growth phase that continued until unfolding was completed according to the nanoDSF profile. This correlation suggests that partial unfolding exposes additional interaction sites on HMGB1, enabling more extensive complexation with heparin or already existing heparin–HMGB1 complexes and promoting the formation of stabilized associates. Indeed, at physiological ionic strength, the electrostatic attraction between positively charged regions of HMGB1 and the highly sulfated, negatively charged heparin chains is partially screened by salt counterions. Consequently, a higher amount of polyanionic heparin is required to compensate for charge screening and lead to distinct-size associates. Together, these results highlight the central role of electrostatic interactions in the formation and stabilization of HMGB1–heparin associates.

**Table 1.** DLS aggregation onset and nanoDSF melting temperatures of HMGB1–PE complexes.

| **eu-HMGB1** |  |  |  |
| --- | --- | --- | --- |
| salt-free |  |  |  |
| PE type | mol% | DLS onset temp (°C) | nanoDSF $T_{m,app}$ (°C) |
| Heparin | 30 | 34 | 40 |
|  | 50 | 36 | 44 |
| lPGS | 30 | 46 | 50 |
|  | 50 | 62 | 52 |
| **bac-HMGB1** |  |  |  |
| salt-free |  |  |  |
| Heparin | 30 | 35 | 41 |
| physiological salt |  |  |  |
| Heparin | 30 | 35 | 45 |
|  | 100 | 37 | 43 |

The absence of association into the bigger complexes at higher linear PE concentrations (Fig. 3a_3_, 3b_3_; Fig. 5b) may be explained by a change in the binding regime. At low PE concentration, multiple HMGB1 molecules can bind to the same polymer chain, thereby increasing the probability of interprotein contacts and promoting the formation of higher-order assemblies (see Scheme 1). These assemblies may be stabilized by a combination of electrostatic interactions, hydrophobic contacts, and partial exposure of oligomerization-prone protein regions during unfolding. In contrast, at higher PE concentrations, the system may approach a regime in which individual HMGB1 molecules are bound to separate polymer chains. This would reduce inter-complex connectivity and thereby suppress the formation of extended associations. A similar concentration-dependent association behavior has been reported for lysozyme/heparin systems, where fibril formation was observed only within a defined range of heparin concentrations [26–28].

**Scheme 1.**
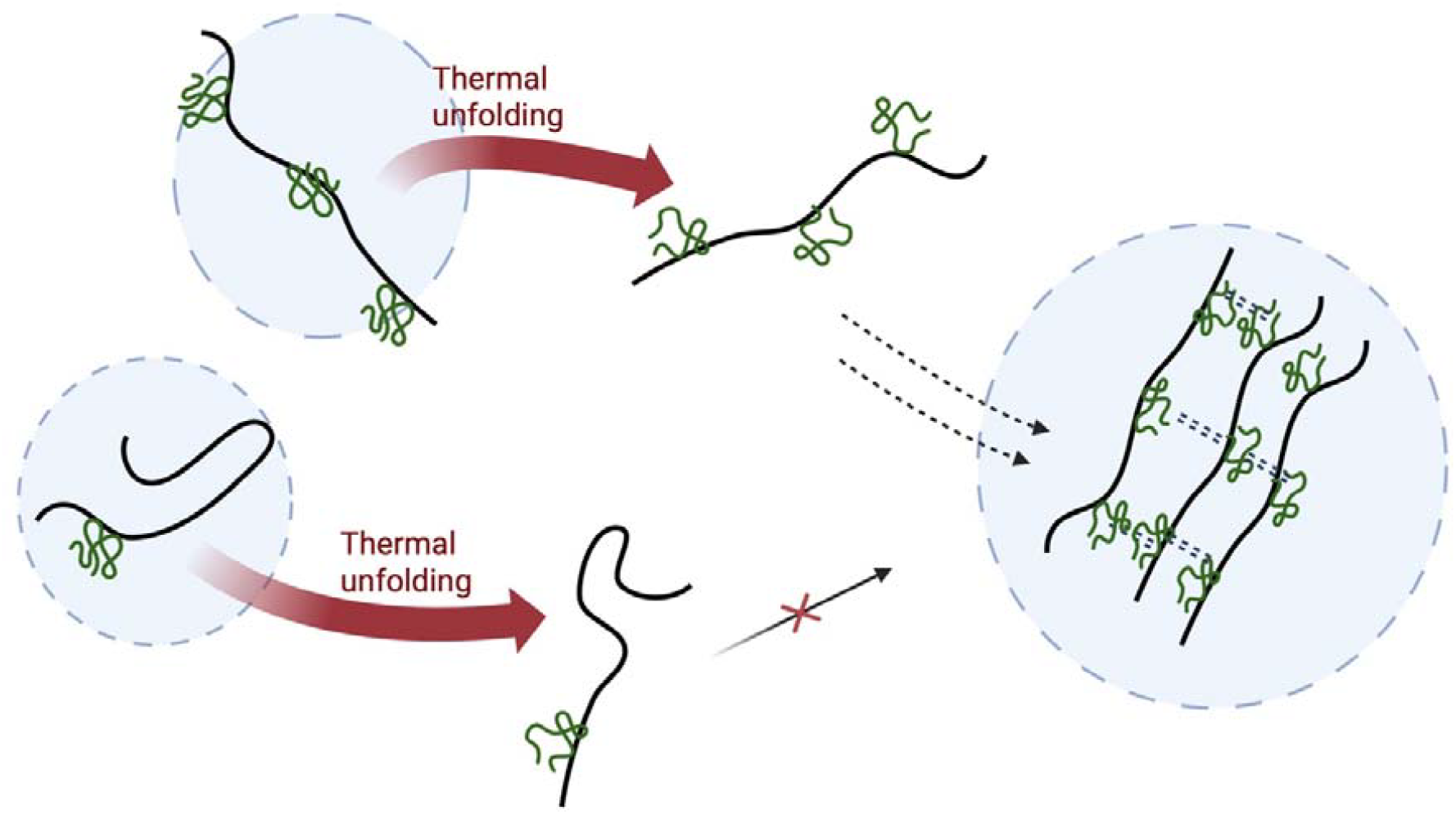
Proposed mechanism of temperature-induced HMGB1–PE complexation. Green coils represent HMGB1 proteins, while black strings represent linear PEs. The blue bubble represents the hydrodynamic shell of the HMGB1–PE associates.

## Conclusion

In this work, we show that bac-HMGB1 as well as eu-HMGB1 interact with linear sulfated polyelectrolytes in a polymer-dependent, glycosylation-independent, but mechanistically related manner. Heparin and lPGS produced similar effects on the thermal response of HMGB1, suppressing the main nanoDSF unfolding transition, improving the protein refolding, and partially preserving the secondary structure during heating. These results indicate that both linear polysulfates stabilize HMGB1 against irreversible thermal unfolding caused by protein oligomerization upon heating, despite their different chemical complexity and backbone flexibility. The comparison with non-sulfated lPGOH confirmed that this effect is not caused by the presence of a polymer alone but requires sulfate-mediated electrostatic interactions. A particularly important finding is the formation of defined HMGB1–polyelectrolyte assemblies ranging in size from 20 to 40 nm upon heating. This behavior occurred within a specific concentration window depending on the polymer flexibility and in the physiologically relevant temperature range. Although the exact nature of these assemblies remains to be clarified, their formation suggests that polyelectrolytes do not simply suppress HMGB1’s extended oligomerization to very large sizes during unfolding, but can redirect the protein into alternative supramolecular states. It was also found that it is possible to modulate this interaction by varying the salt concentration. Here, the results at physiological salt conditions underscore electrostatic interactions as the main driver of the polyelectrolyte dependent association behavior, as it was shown that a higher heparin concentration is needed to achieve controlled association. This previously unrecognized behavior is relevant for understanding how HMGB1 is modulated by charged polymers.

## Acknowledgments

This work was funded by the Deutsche Forschungsgemeinschaft (DFG, German Research Foundation) – 434130070 within the GRK 2662. We thank Prof. Rainer Haag for the support.

## Data availability

Raw and meta data is available under DOI:

