## Supplemental Figure S1 for "Thermodynamic properties and stability of HMGB1 complexes with linear polyelectrolytes elucidated by nano differential scanning fluorimetry"

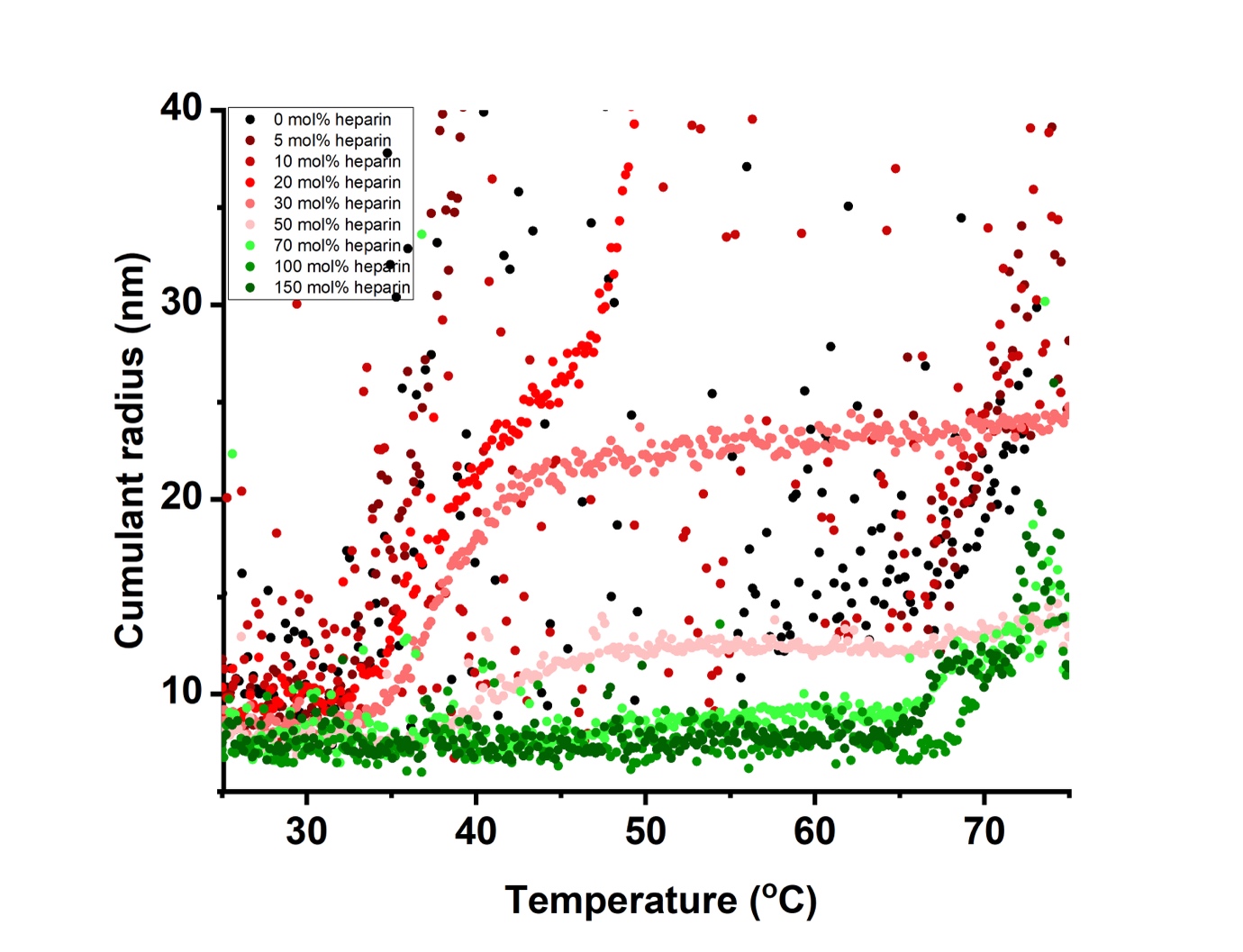

**Fig. S1** Temperature dependent cumulant radius analysis of eu-HMGB1 in the presence of heparin in different molar ratios.
